# Mental Timing in Locomotion: Walking Across Real and Imagined Distances

**DOI:** 10.64898/2026.09.02.748865

**Authors:** Elise E Van Caenegem, Gaëlle Lorge, Marcos Moreno-Verdú, Charlène Truong, Baptiste M Waltzing, Robert M Hardwick

## Abstract

The present study aimed to directly replicate the seminal findings of Decety et al. (1989), while also extending the original work with a larger sample size and additional statistical analyses.

Thirty healthy adults completed both imagined and executed walking tasks over distances of 5, 10, and 15 meters. Participants also completed the Movement Imagery Questionnaire-3 (MIQ-3). Temporal correspondence between imagined and executed walking was assessed using repeated-measures ANOVA, correlation analyses, Bayesian equivalence tests, and Bland–Altman analyses.

Walking duration increased significantly with distance, while no significant differences were observed between imagined and executed walking times. Bayesian equivalence tests provided evidence supporting temporal equivalence between the two modalities, and Bland–Altman analyses indicated good agreement with minimal systematic bias. Strong positive correlations were found between imagined and executed durations at all distances. MIQ-3 scores were not significantly associated with differences between imagined and executed walking durations.

Overall, the findings successfully replicate and extend the original results reported by Decety et al. (1989), providing additional evidence supporting temporal equivalence between motor imagery and movement execution. By combining a larger sample size with contemporary statistical approaches, the present study strengthens support for the principle of functional equivalence for highly automated and cyclic movements.

## 1. Introduction

### 1.1 Theoretical background

Motor imagery refers to the process of mentally simulating an action without actually performing the movement. One of the classic empirical approaches to assessing the quality of this simulation is based on mental chronometry (Posner, 1978), which refers to the time course of information processing by the nervous system. In the case of motor imagery, this involves comparing the time taken to perform an action with the time taken to imagine that same action. This concept in motor imagery was first developed by Decety et al. (1989). Since then, a great amount of research has been carried out to evaluate the temporal fidelity of motor imagery, as an indication of its validity and usefulness in contexts as varied as sport, rehabilitation and cognitive neuroscience (for reviews see Guillot & Collet, 2005; Guillot et al., 2012).

Mental chronometry research is based on empirical evidence showing that the duration of imagined movements closely matches the duration of its execution (Decety et al., 1989). This later inspired the formalization of the motor simulation theory (Jeannerod, 1994). Based on the concept of ‘functional equivalence’, motor simulation theory proposes that motor imagery recruits sensorimotor processes like those activated during actual execution. This hypothesis is supported by a meta-analysis from Hardwick et al. (2018), which shows that motor imagery and movement execution share a cortico-subcortical network comprising premotor cortex, parietal cortex, cerebellum, and basal ganglia. However, although motor imagery and movement execution engage overlapping neural substrates, the extent and pattern of neural activation are not fully equivalent, suggesting that functional equivalence may be more nuanced than initially proposed (Van Caenegem et al., 2024).

Several studies have examined the temporal equivalence of actions across various paradigms. For example, Decety et al. (1989) showed that imagined and real walking durations did not differ significantly, and that the timing increased in proportion to the distance to be covered in both modalities, reflecting similar temporal dynamics during real and imagined walking. Similar results were obtained in pointing tasks (Gentili et al., 2006), writing tasks (Decety & Michel, 1989), and arm movements with load manipulation (Papaxanthis et al., 2002), where imagined durations varied according to the mechanical complexity of the action to be simulated. More recent studies have provided further evidence that task-related constraints are also reflected during motor imagery, including constraints related to movement coordination (Dahm & Rieger, 2016) and movement difficulty as described by Fitts’ law (Czilczer et al., 2026). These data indicate that motor simulation integrates not only the kinematics, but also the dynamic constraints of the movement.

However, the correspondence between executed and imagined durations is not always perfect, and several factors can affect this synchrony. The difficulty of the task (Jeannerod, 1995, 1999), prior experience of the action to be simulated (Guillot et al., 2012), age (Skoura et al., 2005), the type of instruction given (internal vs. external, kinesthetic vs. visual), or the amount of physical practice (McAteer et al., 2025) can influence the temporal accuracy of motor imagery (Callow & Hardy, 2004). In addition, competing cognitive tasks can interfere with motor simulation and alter temporal accuracy; Glover and Baran (2017) showed that the addition of a secondary verbal working memory task significantly disrupted the timing of motor imagery, suggesting that it actively mobilises attentional and executive resources. This last finding somewhat challenges the principle of functional equivalence, suggesting that motor imagery may rely more on cognitive processes than movement execution. Further evidence to support this proposal is provided by the recent meta-analytic work by Van Caenegem et al. (2026), which showed that motor imagery shares more neural activity with working memory than with movement execution.

### 1.2 Aim of the study

The aim of this study was to replicate and extend the protocol established by Decety et al. (1989), which consisted of comparing imagined and executed walking timing across different distances. This temporal correspondence is a key prediction of the Motor Simulation Theory (Jeannerod, 2001), which argues that motor imagery relies on processes similar to those involved in motor execution. Replicating this finding with contemporary methods would therefore strengthen its support for automatic movement such as walking.

While this seminal study was the first to examine the use of mental chronometry in motor imagery research, certain methodological limitations should be acknowledged. First, only 10 participants completed the imagery tasks in the original study. This is a rather limited sample size in the contemporary context, where larger samples are generally expected to improve the reliability and external validity of findings (Button et al., 2013). Secondly, there was limited evidence of direct statistical equivalence between the durations of imagined and executed walking. Direct comparisons of the durations of imagined and executed walking were made via paired-samples t-tests which found non-significant differences; by contrast, modern approaches have introduced techniques to confirm statistical equivalence between measurements. Similarly, while very strong correlations (i.e. r = 0.89 - 0.99) were reported, the analyses appeared to use a mixture of within- and between-participant measures, which can inflate r values and increase the likelihood of false positives (Ranganathan & Aggarwal, 2016). Finally, while all participants were indicated to be “good imagers” based on questionnaires, no individual scores were presented for the participants, and comparisons between motor imagery ability and the temporal precision of imagined actions were not presented. In the present study, we will attempt to overcome these limitations via a direct methodological replication with analytical extension. This will primarily be achieved by increasing the number of participants and conducting statistical analyses using modern approaches that can allow both the confirmation and the rejection of the null hypothesis (e.g. Bayesian analyses, equivalence tests, etc). Supplementary analysis will examine possible relationships between imagery ability (as defined via questionnaire scores) and the temporal precision of imagined actions. Importantly, we note that even in the context of the limitations mentioned above, we expect to observe a strong temporal correspondence between executed and imagined timing.

## 2. Methods

### 2.1 Sample Size Determination

Using a Bayesian sequential sampling design, data collection continued until a minimum sample size of n = 30 was reached. Thereafter, our plan was to apply a stopping rule such that data collection would cease once we reached a Bayes Factor (BF) ≥ 3 in favour of either the null or alternative hypothesis (indicative of ‘moderate’ evidence according to established benchmarks (Andraszewicz et al., 2015; Jeffreys, 1998)) for the comparisons between the durations of imagined and executed actions at each of the three distances examined.

### 2.2 Participants

Thirty healthy participants were recruited to take part in this study. Participants were recruited from UCLouvain student community in Belgium. The participants were 12 males and 18 females with an average age of 21.9 ± 1.7 years (mean ± SD). Of the 30 participants, only 28 were right-handed, and 2 were left-handed. Participants were asked if they had any previous experience of motor imagery; only 2 participants mentioned having used motor imagery before, and all other participants reported they were naïve to previous imagery use. Participants were financially compensated (10€) for their time if they fully completed the study (full completion was approximately 45 minutes). All participants provided informed consent prior to participation. They all completed the study and no participant was excluded. This experiment received ethical approval from the UCLouvain Psychological Sciences Research Ethics Committee (reference: Project 2024–79-Bis, Date: 06/12/2024).

### 2.3 General procedure

#### 2.3.1 Movement imagery questionnaire 3 (MIQ-3)

Participants were firstly asked to complete the French version of the MIQ-3 (Robin et al., 2021; for the English version see Williams et al., 2012) in order to assess participants’ ability to generate mental images of movement in their mind’s eye. Each item required participants to physically perform one of four movements, then to imagine doing the same movement using either kinesthetic imagery (imagining the sensations associated with the movement), internal visual imagery (seeing the action from a first-person perspective), or external visual imagery (viewing the action from a third-person perspective). Afterward, for each item, participants rated how easily they could feel or visualize the imagined movement on a seven-point scale from 1 (very difficult to feel/see) to 7 (very easy to feel/see).

#### 2.3.2 Description of the task and experimental setup

In order to be able to compare the results obtained with those of Decety et al. (1989), we reproduced the task procedure as closely as possible based on the methods reported by their manuscript. A walking path was created using orange tape lines, which were placed on concrete at distances of 0m, at 5m, 10m and 15m.

At the start of the experiment participants received the following instructions for the imagery task: “With your eyes closed, after having fixed the position of the tape line corresponding to the distance in your mind, you have to imagine yourself walking the distance indicated, kinesthetically and visually in the first-person perspective. Try to keep your speed constant and similar to the speed you would have adopted when walking in real life. You are not allowed to count in your head.”

Prior to each trial the experimenter instructed the participant about the required modality (i.e. executed or imagined walking) and the distance (i.e. 5, 10, or 15m). Participants held a digital memory stopwatch (500-Memory DT500 Stopwatch USB version Ref:024111) in their dominant hand and were instructed to activate it when initiating (or imagined initiating) the first step, and to stop it when they crossed (or imagined crossing) the corresponding tape line on the floor. The duration of each trial was recorded in the digital memory of the stopwatch, and thus the participant and the experimenter were blind to the times recorded throughout the experiment.

Ten trials of each modality for each distance were completed in a different random order for each participant to avoid block effects. As such a total of 60 trials were completed (2 modalities x 3 distances x 10 trials). The experiment was carried out in a double-blinded design. Neither the experimenter (GL) nor the participants were aware of the experiment’s exact purpose or expected results.

### 2.4 Data analysis

All statistical analyses were performed with JASP Version 0.19 (for frequentist, Bayesian, and equivalence tests) and with R version 4.4.1 for visualization.

Replicating the procedure of Decety et al. (1989), a 2×3 repeated-measures ANOVA was performed to assess the main effects of modality (execution vs. imagery) and distance (5m, 10m, or 15m), as well as their interaction. Effect sizes are reported as eta squared (η², not partial eta squared) and were interpreted according to Cohen’s (1988) benchmarks, with η² = .01, .06, and .14 indicating small, medium, and large effects, respectively. Extending this procedure to assess not only possible differences between modalities, but also whether their durations could be considered practically similar, Equivalence Bayesian Paired Samples T-tests were conducted for each distance using a default prior distribution (Cauchy distribution with a location parameter of 0 and a scale parameter of 0.707), corresponding to a standardized region of practical equivalence (ROPE) (Kruschke, 2018) with predefined regions of practical equivalence δ ∈ [-0.1, 0.1]. The resulting Bayes Factors (BF) were interpreted according to standard thresholds with inconclusive evidence (=1), anecdotal (1–3), moderate (3–10), strong (10–30), very strong (30–100), or extreme (>100) evidence (Kass & Raftery, 1995). In line with Decety et al. (1989), Pearson correlations were used to assess the strength of association between executed and imagined times at each distance with negligible (<0.1), weak (0.1–0.4), moderate (0.4–0.7), strong (0.7–0.9), or very strong (>0.9) (Schober et al., 2018). Further analysis examined possible correlations between MIQ-3 scores and the temporal difference (i.e. duration of imagery–execution) for the different walking distances. Data used for correlation analyses were screened for outliers using the “robust correlation” MATLAB toolbox (Pernet et al., 2013). Agreement between execution and imagery was further explored using Bland-Altman plots, which visualized systematic biases and the percentage of data within limits of agreement (Bland & Altman, 1999).

## 3. Results

### 3.1 Repeated measures ANOVA 2x3

Following Decety’s original analysis, a 2x3 repeated measures ANOVA was performed to observe whether there was an effect of distance (5m, 10m, 15m), modality (execution or imagery) or whether there was an interaction between distance and modality (Fig. 1). Because the assumption of sphericity was violated for Distance, Greenhouse–Geisser corrections were applied when appropriate. With regard to distance, we observed a significant main effect with a large effect size (F(1.10, 31.81) = 645.49, p < .001, η² = .889), whereby imagined and executed times both increased as walking distance increased. Post hoc comparisons showed that all distances significantly differed from one another: times were higher at 10 m than at 5 m (ΔM = 3.415, SE = 0.134, *t*(29) = 25.487, *p* < .001), higher at 15 m than at 5 m (ΔM = 6.778, SE = 0.260, *t*(29) = 26.026, *p* < .001), and higher at 15 m than at 10 m (ΔM = 3.363, SE = 0.145, *t*(29) = 23.217, *p* < .001). There was no significant main effect of modality (F(1, 29) = 0.67, p = .419, η² = .001), nor a significant interaction between distance and modality (F(1.27, 36.94) = 0.05, p = .877, η² < .001).

**Figure 1:**
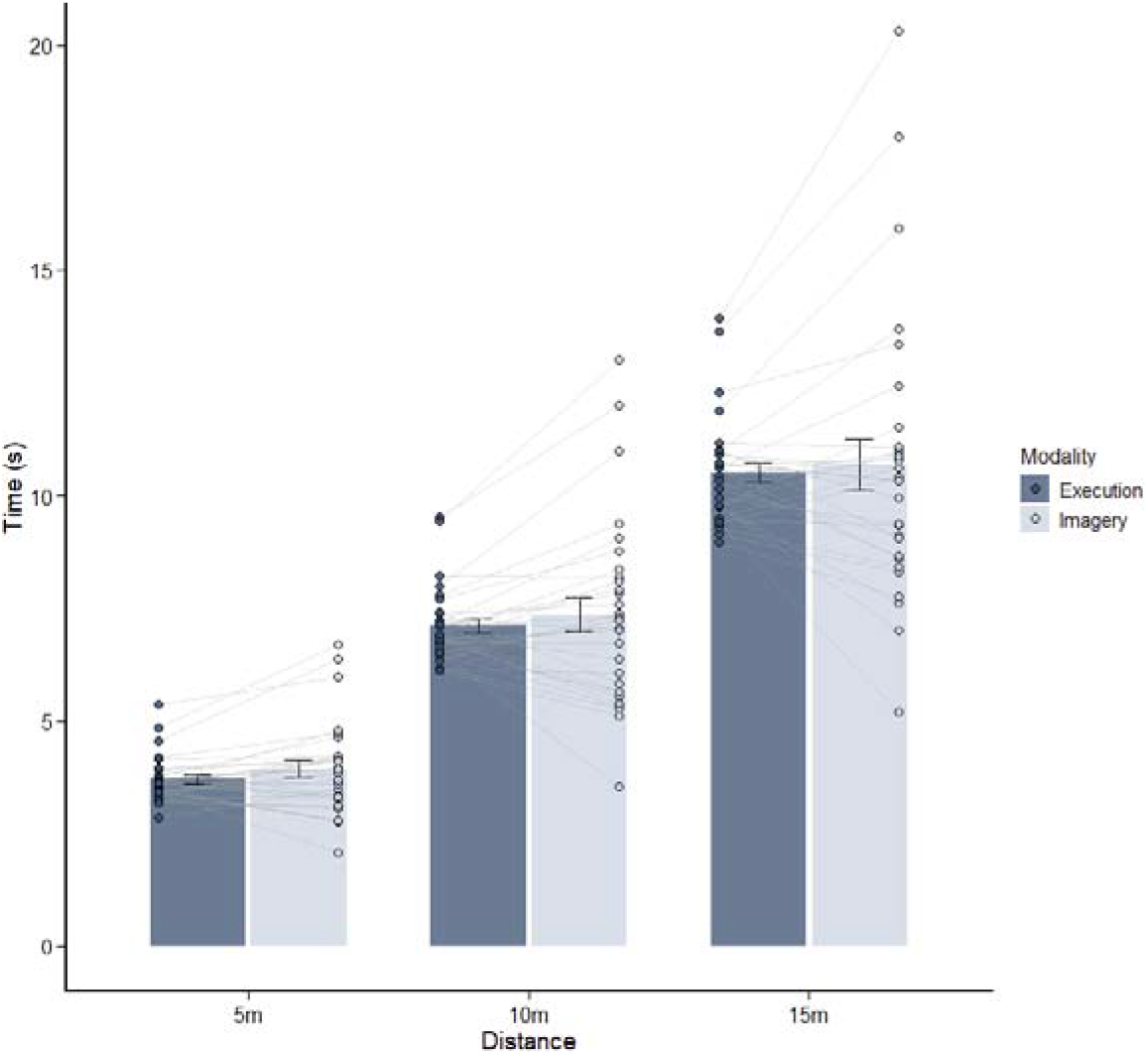
Walk timing mean across all participants for both modalities and for each distance. Individual points represent each participant for each modality and error bars present standard error of the mean.

The mean and standard deviations (SD) for executed and imagined timing for all distances are shown in the table in supplementary materials (Table S1).

### 3.2 Equivalence tests

Bayesian paired-samples T-tests were used to assess evidence for equivalence between executed (ME) and imagined (MI) walking durations at each distance. Across all tests, the Bayes Factor for the overlap hypothesis (BF^NOH^ _01_) indicated that the data was ≥ 3.136 times more likely to support the hypothesis of equivalence, and the Bayes Factor for the non-overlap hypothesis (BF^NOH^ _01_) indicated the data was ≥ 3.969 times more likely to lie in the region of equivalence than the region of non-equivalence. Full values for each test are presented in Table 1.

**Table 1:** Equivalence Bayesian Paired Samples T-Test. For model comparisons, “δ ∈ I vs. H” refers to evidence in support of the hypothesis of equivalence (BF^NOH^ _01_), while “δ ∈ I vs.δ ∉ I” refers to evidence in support of the non-overlap hypothesis (BF^NOH^_01_).

|  |  |  | Model Comparison | BF | error % |
| --- | --- | --- | --- | --- | --- |
| ME_5m | - | MI_5m | $\delta \in I$ vs. $H\Box$ | 3.136 | $1.446 \times 10^{-5}$ |
| | | | $\delta \in I$ vs. $\delta \notin I$ | 3.969 | $2.285 \times 10^{-5}$ |
| ME 10m | - | MI 10m | $\delta \in I$ vs. $H\Box$ | 4.248 | $2.758 \times 10^{-5}$ |
| | | | $\delta \in I$ vs. $\delta \notin I$ | 6.238 | $3.756 \times 10^{-5}$ |
| ME_15m | - | MI_15m | $\delta \in I$ vs. $H\Box$ | 4.746 | $1.997 \times 10^{-5}$ |
| | | | $\delta \in I$ vs. $\delta \notin I$ | 7.510 | $2.524 \times 10^{-5}$ |

### 3.3 Correlations

Replicating Decety et al. (1989) analysis, Pearson correlations between executed (ME) and imagined (MI) times were conducted. Strong and significant correlations were observed for the corresponding distances (Table 2), indicating a strong linear relationship between the executed and imagined walking times (Fig. 2).

**Figure 2:**
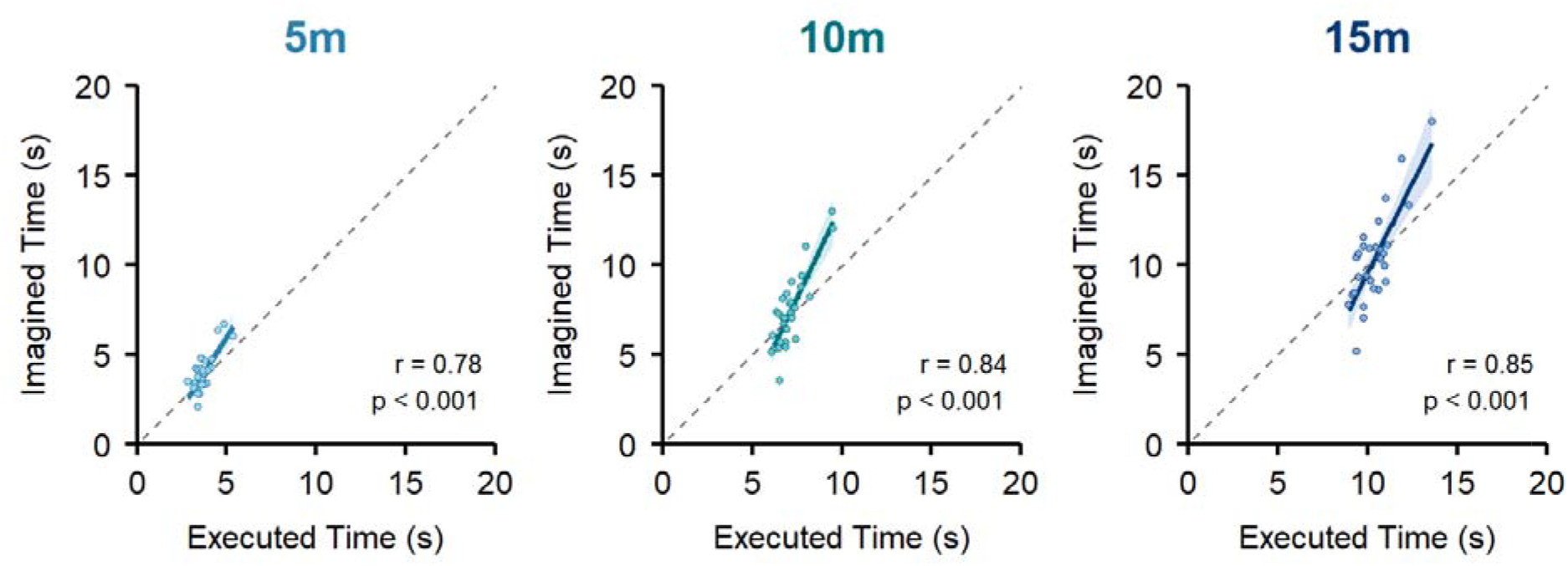
Correlations between executed and imagined walking timing across all distances.

**Table 2:** Pearson’s Correlations.

| Variable |  | ME_5m | MI_5m | ME_10m | MI_10m | ME_15m | MI_15m |
| --- | --- | --- | --- | --- | --- | --- | --- |
| 1. ME_5m | Pearson's r | — |  |  |  |  |  |
| 2. MI_5m | Pearson's r | **.781** | — |  |  |  |  |
|  | p | **< .001** |  |  |  |  |  |
|  | 95% CI | **[.586, .891]** |  |  |  |  |  |
| 3. ME_10m | Pearson's r | .937 | .766 | — |  |  |  |
|  | p | < .001 | < .001 |  |  |  |  |
|  | 95% CI | [.870, .970] | [.560, .883] |  |  |  |  |
| 4. MI_10m | Pearson's r | .842 | .932 | **.841** | — |  |  |
|  | p | < .001 | < .001 | **< .001** |  |  |  |
|  | 95% CI | [.691, .922] | [.861, .967] | **[.689, .922]** |  |  |  |
| 5. ME_15m | Pearson's r | .906 | .740 | .982 | .814 | — |  |
|  | p | < .001 | < .001 | < .001 | < .001 |  |  |
|  | 95% CI | [.809, .954] | [.517, .869] | [.962, .991] | [.643, .908] |  |  |
| 6. MI_15m | Pearson's r | .836 | .927 | .865 | .961 | **.853** | — |
|  | p | < .001 | < .001 | < .001 | < .001 | **< .001** |  |
|  | 95% CI | [.681, .919] | [.851, .965] | [.734, .934] | [.919, .982] | **[.712, .928]** |  |

Note. Values in bold are the executed–imagined comparisons at the same distance, i.e. the correlations plotted in Fig. 2.

Executed and imagined walking times were strongly correlated at 5 m, r(28) = .78, 95% CI [.59, .89], p < .001, at 10 m, r(28) = .84, 95% CI [.69, .92], p < .001, and at 15 m, r(28) = .85, 95% CI [.71, .93], p < .001. These results indicate that participants who were relatively slower in the executed walking condition also tended to be relatively slower when imagining the same distance.

However, if we look at the other values, strong and significant correlations were also observed between the different distances, within both modalities. For example, executed walking times at 5 m and 10 m were very strongly correlated (r(28) = .94, 95% CI [.87, .97], p < .001), as were imagined walking times at 5 m and 10 m (r(28) = .93, 95% CI [.86, .97], p < .001). This suggests that the participants were consistent in their relative timing across distances, i.e. participants who were relatively slower (or faster) at one distance tended to remain relatively slower (or faster) at the other distances (Fig. 3).

**Figure 3:**
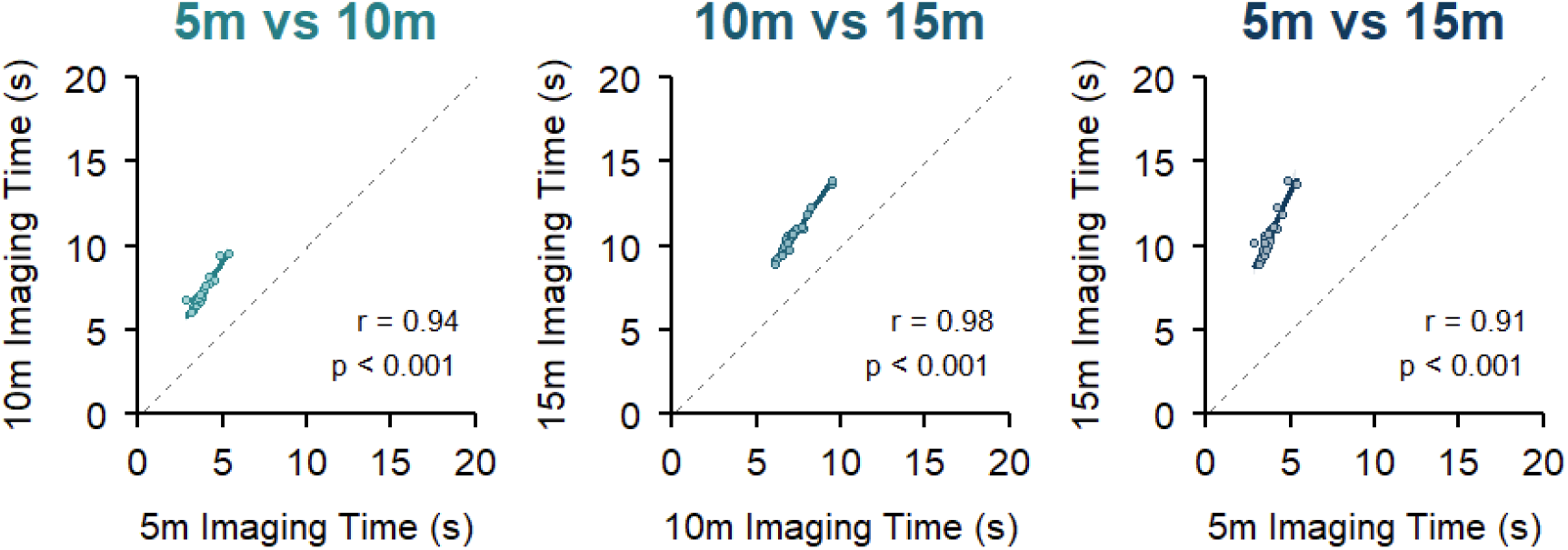
Correlations between imagery timings for each distance.

### 3.4 Bland-Altman test

To assess whether the timing of imagined and executed movements showed acceptable agreement, a Bland-Altman test was carried out. For each distance, we calculated the limits of agreement, corresponding to ± 1.96 multiplied by the SD of the mean difference between imagined and executed timings (dotted line in Fig. 4). All the points were then averaged (full line in Fig. 4). A mean difference close to zero indicates that there is no systematic bias between imagined and actual timing.

**Figure 4:**
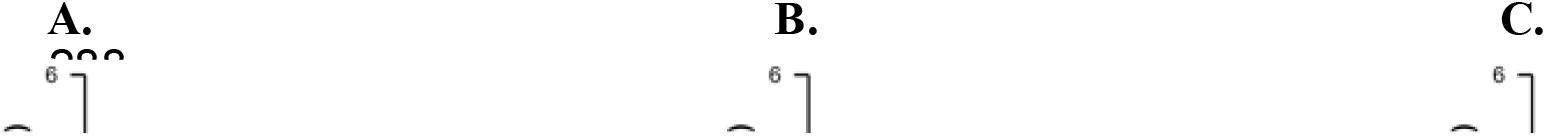
**A.** Bland-Altman plot for 5m. **B.** Bland-Altman plot for 10m. **C.** Bland-Altman plot for 15m.

The results for the upper and lower limits and the average points showed narrow limits of agreement for 5m (Mean = -0.22 seconds [-1.61, 1.17]), 10m (Mean = -0.24 seconds [-3.09, 2.61]) and 15m (Mean = -0.17 seconds [-4.49, 4.15]).

### 3.5 MIQ-3

The mean MIQ-3 score across all participants was 60.16 (SD ± 7.97). The correlation between the absolute time difference between executed/imagined movements and MIQ-3 global scores (Fig. 5) was not significant (r(28) = -0.23, 95% CI [-.55, .14], p = 0.21).

**Figure 5:**
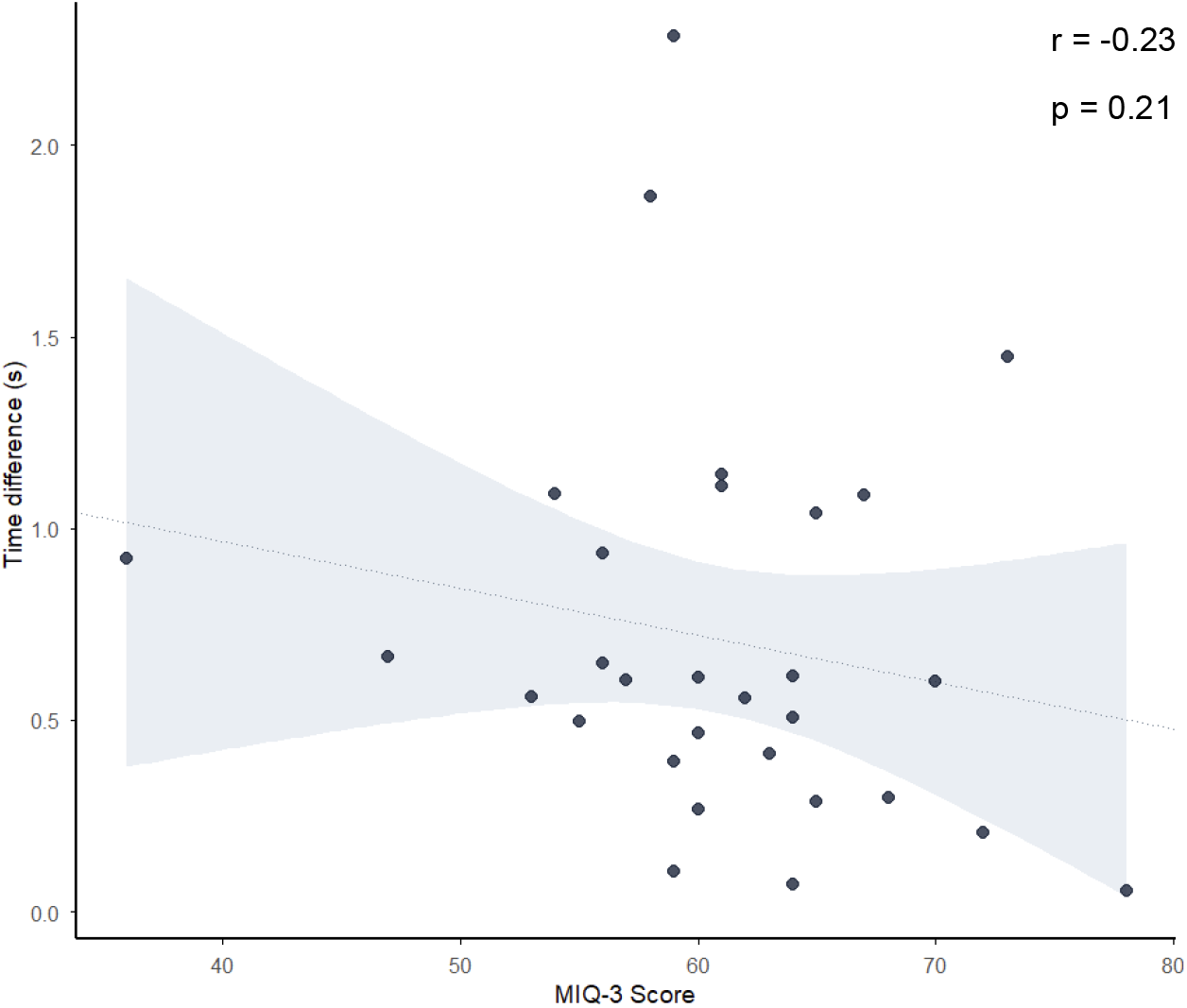
Correlation between timing differences and MIQ-3 global score.

Correlations for sub-scores can be found in the supplementary materials (Table S3).

## 4. Discussion

### 4.1 General Overview

The aim of this study was to replicate and extend the protocol established by Decety et al. (1989), while addressing methodological limitations in the original work. Our results confirmed a strong temporal correspondence between imagined and actual walking, with no significant differences between these modalities. Further analysis identified strong correlations (i.e. r ≥ 0.78) for imagined and executed walking times at each distance. These results replicate those observed in the original study (Decety et al., 1989), but with greater robustness given the larger sample size and more advanced statistical tests. These results show that participants are consistent across modalities. A participant who walks slowly will tend to imagine walking slowly as well. The results also show that participants are consistent across distances. A participant who walks slowly over 5 metres will do so over 10 and 15 metres as well.

The main finding of this study was to empirically demonstrate that the time taken to walk or imagine walking the same distance is statistically equivalent. The combined use of an ANOVA, a Bayesian equivalence test, and the Bland-Altman plot allowed us to go beyond simply finding no significant difference between the measures, and to instead conclude that there is true equivalence between the imagined and actual walking times.

### 4.2 Equivalence between imagined and physical timings

Based on these results presented above, the data support the motor simulation theory’s functional equivalence hypothesis (Jeannerod, 1994), suggesting that the temporality of motor imagery and real movement are equivalent. Our findings are consistent with other studies that have examined the temporal components of motor imagery and execution. Temporal similarities have been observed in studies focusing on walking (Courtine et al., 2004), but also in other tasks such as writing (Decety & Michel, 1989), pointing tasks (Gueugneau & Papaxanthis, 2010), and reaching and grasping tasks (Errante et al., 2019). Although these studies did not show any significant differences between executed and imagined timings, none of them carried out equivalence tests, unlike our study.

However, not all studies show similarities between actual and imagined movement times. According to Guillot and Collet (2005), timings are typically identical for highly automatic movements (e.g. over-practiced movements such as pointing or reaching to grasp objects) and cyclic movements (such as walking, rowing, running) but differ for more complex tasks. Imagined movement times may be underestimated (faster) when imagery is performed in a stressful context such as before a competition (Calmels & Fournier, 2001; Munroe et al., 2000), when the entire movement is not imagined but only a sequence (Vieilledent, 1996), or when the task involves lengthy and complex preparation (Deschaumes-Molinaro et al., 1992). Conversely, in some cases imagined movement times may be overestimated (longer). This is particularly the case for technically complex tasks, and this discrepancy increases as the task becomes more complex (Cerritelli et al., 2000; Georgopoulos et al., 1989), but also when the movement to be imagined is fast and requires intense concentration (Coello & Orliaguet, 1992).

In this sense, our results correspond to the profile of tasks for which equivalence is observed most consistently. For healthy adults walking is a highly practised, automated, cyclic motor pattern thanks to the experience acquired throughout life. This facilitates a temporal correspondence between imagination and execution due to a high level of familiarity with the task.

### 4.3 No significant correlation between MIQ-3 results and imagery timing

Our results did not demonstrate a significant correlation between task performance and the results of a questionnaire designed to measure motor imagery ability (i.e. the MIQ-3). While this could be due to the relatively small sample size for correlational analyses in the present study, we note that previous research that has been adequately powered to detect even small correlations found similar results. In a sample of 198 participants, Williams et al. (2015) found no correlation between the score at MIQ-3 and the time taken to complete or imagine the various items in this questionnaire. The fact that no correlation was observed can potentially be explained by the fact that these assessments examine different aspects of motor imagery; that questionnaires can be used to assess the vividness of imagery (i.e. the ability to generate an image), while chronometry assesses the ability to maintain that image (for similar results and discussion of this topic see Moreno-Verdú et al., 2026).

### 4.4 Strengths and limitations

Our paper replicated a previous study with several improvements in the methodology to strengthen its conclusion. First, the sample size was tripled, from 10 participants in the original study to 30 participants in ours. In addition to using frequentist statistics, we also used a Bayesian approach, allowing us to examine evidence both for and against the null hypothesis, and thus provide direct evidence for equivalence between the modalities. Finally, we did not simply observe whether a difference existed; we provided converging evidence supporting equivalence through several tests.

However, this study has some limitations. We only used chronometric data to assess temporal equivalence. No neurophysiological data (e.g., EEG) was collected to reinforce this idea of equivalence at the neural level. Furthermore, the MIQ-3 results did not establish a correlation with performance. While this is consistent with previous suitably powered studies (i.e. Moreno-Verdú et al., 2026; Williams et al., 2015), either collecting more participants to confirm this in the present study, or using another test to assess the ability to imagine that required fewer participants to show a correlation could have been implemented. Finally, as is the case for all research examining covert mental processes, it is not possible to verify with complete certainty that participants indeed engaged in motor imagery; while manipulation checks (i.e. different distances) indicate compliance with the task, there was no objective way to verify that participants did not use a strategy to achieve the same timing, even though they were not aware of the expected results of this study.

### 4.5 Social Impact and Responsibility

Motor imagery is widely used in sport, rehabilitation, and neuroscience research. By strengthening confidence in the temporal correspondence between imagined and executed walking, the present replication contributes to the reliability of a finding that is frequently used to justify imagery-based interventions. At the same time, the study highlights that temporal equivalence appears particularly robust for highly automated actions such as walking; as such, researchers should consider these results in this context, and findings should not be generalized indiscriminately to all forms of motor imagery.

## 5. Conclusion

In conclusion, the present study successfully replicated the findings of Decety et al. (1989), demonstrating a strong temporal correspondence between imagined and executed walking durations across different distances. Beyond reproducing the original findings, the use of Bayesian equivalence testing and Bland–Altman analyses provided formal evidence supporting temporal equivalence between imagined and actual walking. These findings strengthen support for the functional equivalence hypothesis in cyclic and highly automated tasks.

## Transparency Statement

We report how we determined our sample size, all data exclusions (if any), all manipulations, and all measures in the study. No participants were excluded from the reported analyses. The authors have no personal, professional, or collaborative connection with the authors of the original study.

## Data, scripts, code, and supplementary information availability

This study was not preregistered. No preregistered protocol or analysis plan was registered prior to data collection. Data, analysis scripts, study materials, and reproducibility instructions are publicly available at: https://github.com/baslaboratory/Walking-timing.git

## Use of Artificial Intelligence Tools

ChatGPT (GPT-5, GPT-5.x, OpenAI) was used to assist with language editing, text revision, and coding support (R). All scientific content, analyses, and interpretations were reviewed and approved by the authors.

## File-Drawer Statement

The present paper reports all studies, analyses, and outcomes conducted by the authors on this topic that are relevant to the research questions addressed in this manuscript. No additional completed studies with unreported results are known to exist.

## Author Roles (CRediT)

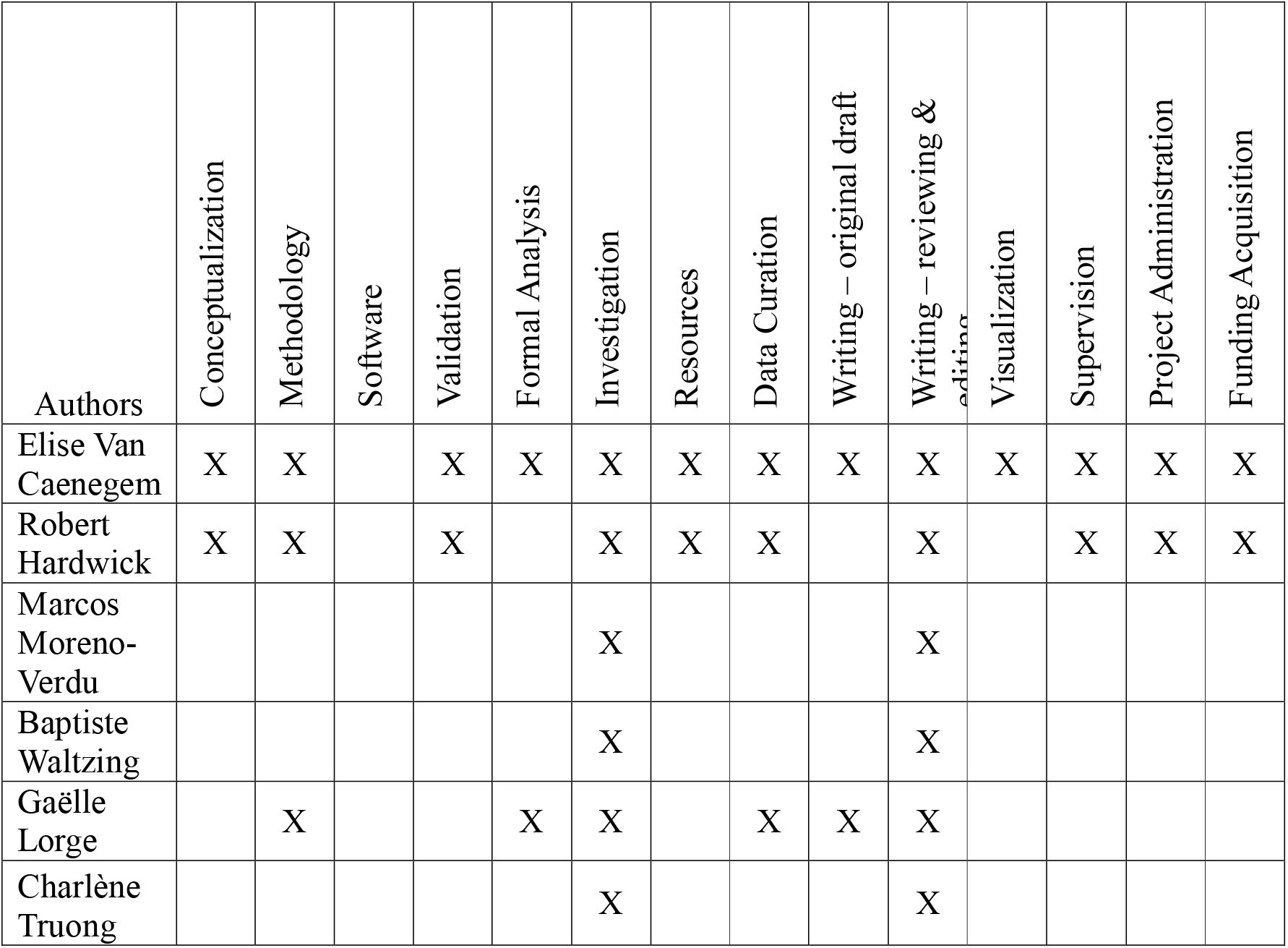

| Authors | Conceptualization | Methodology | Software | Validation | Formal Analysis | Investigation | Resources | Data Curation | Writing – original draft | Writing – reviewing & editing | Visualization | Supervision | Project Administration | Funding Acquisition |
| --- | --- | --- | --- | --- | --- | --- | --- | --- | --- | --- | --- | --- | --- | --- |
| Elise Van Caenegem | X | X |  | X | X | X | X | X | X | X | X | X | X | X |
| Robert Hardwick | X | X |  | X |  | X | X | X |  | X |  | X | X | X |
| Marcos Moreno-Verdu |  |  |  |  |  | X |  |  |  | X |  |  |  |  |
| Baptiste Waltzing |  |  |  |  |  | X |  |  |  | X |  |  |  |  |
| Gaëlle Lorge |  | X |  |  | X | X |  | X | X | X |  |  |  |  |
| Charlène Truong |  |  |  |  |  | X |  |  |  | X |  |  |  |  |

## 6. Supplementary materials

**Table S1:**
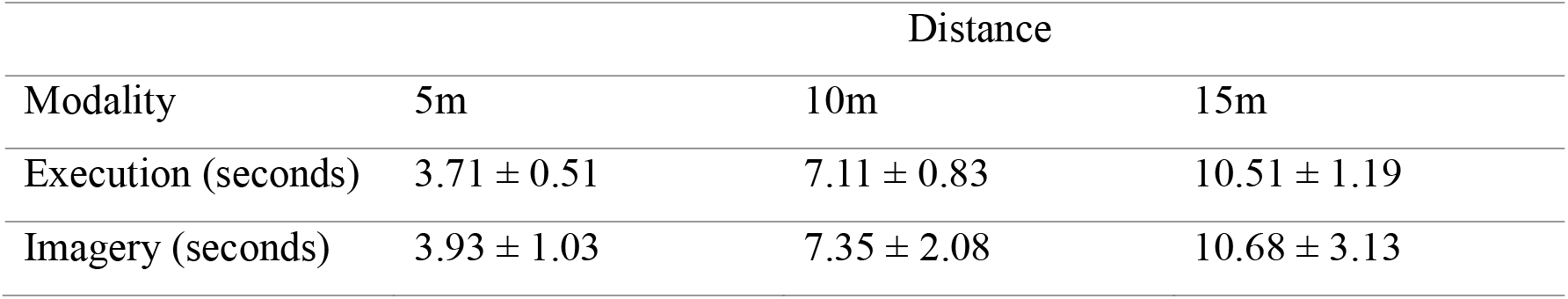
Mean and standard deviations in seconds.

**Table S2:**
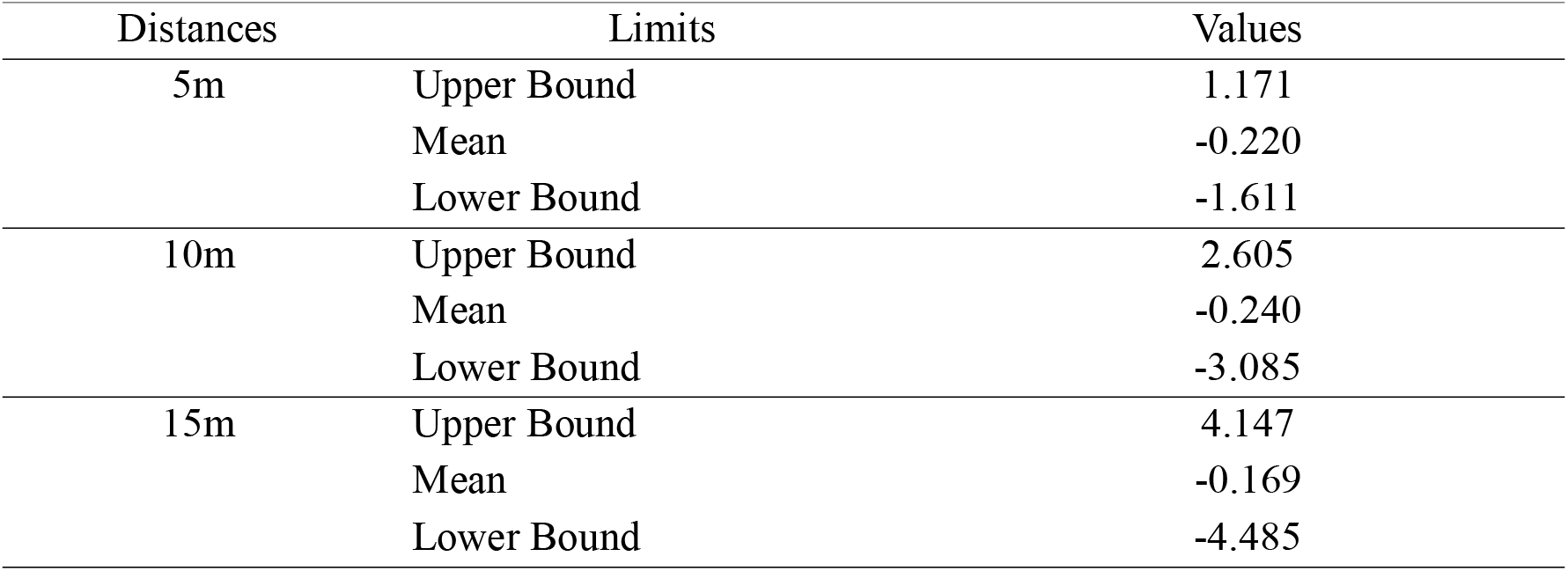
Bland-Altman Test.

**Table S3:**
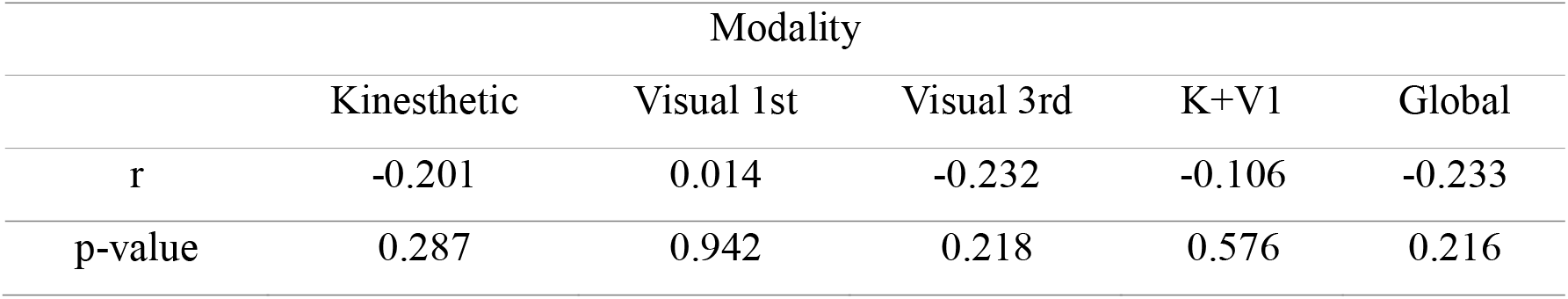
Correlations with MIQ-3 sub-scores.

**Figure S1:**
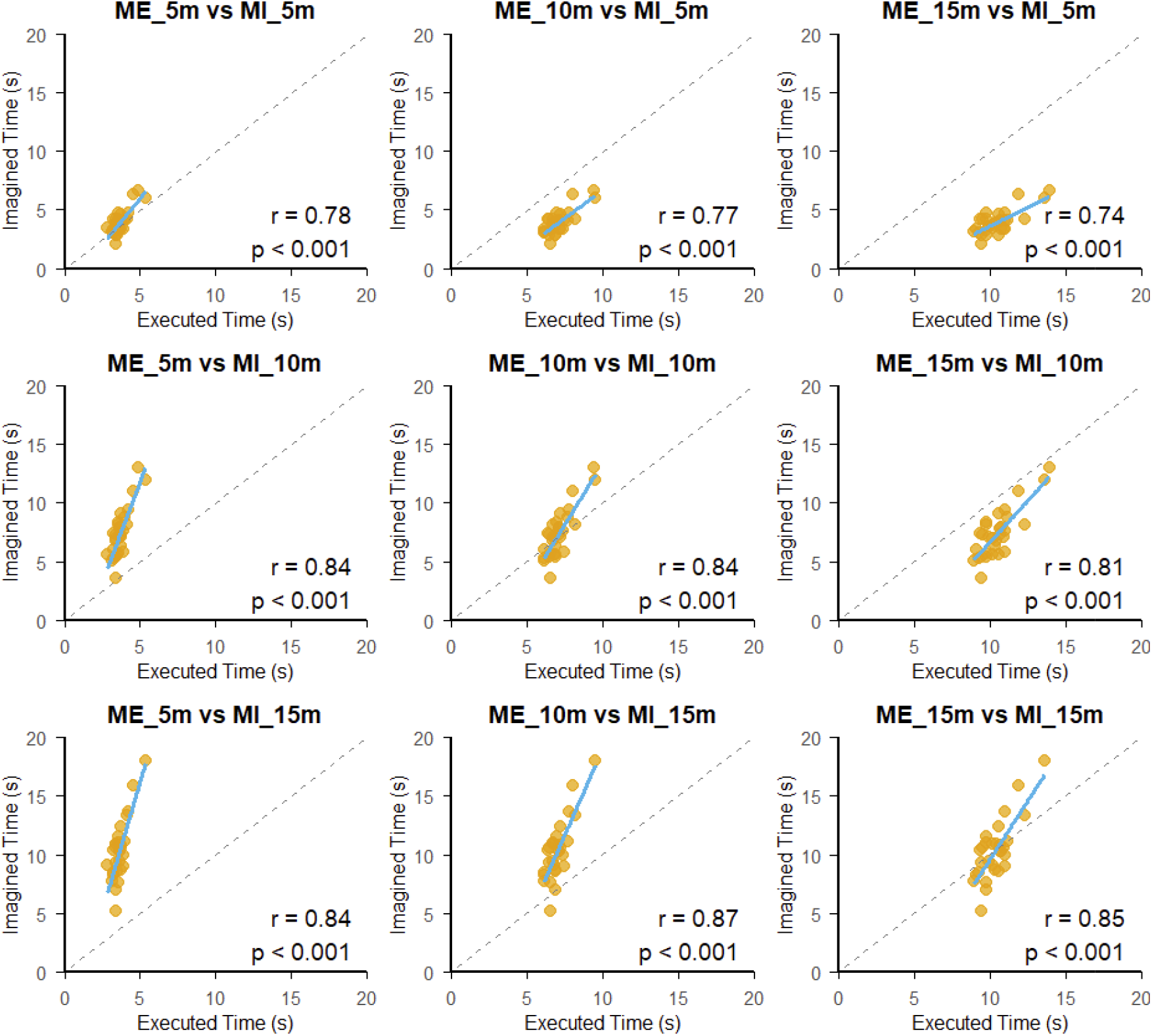
Correlations across all distances and all modalities.

**Figure S2:**
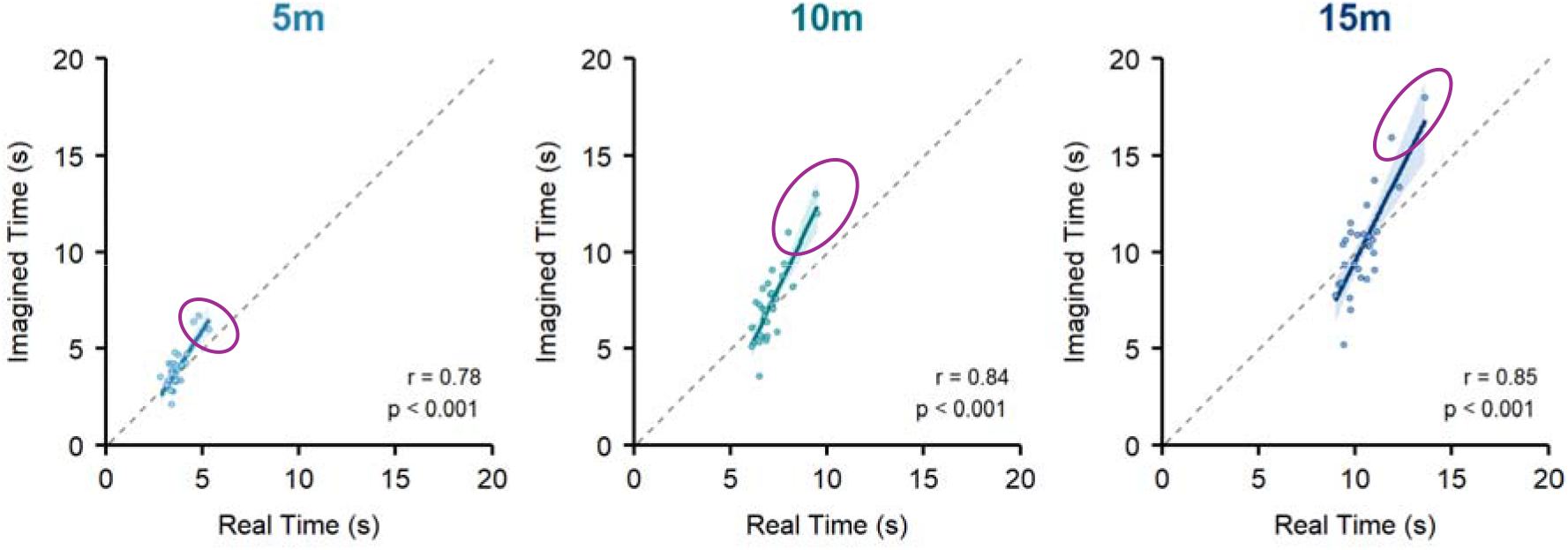
Robust correlations. The points circled in purple are the outliers detected by robust correlation analysis. By removing these points, coefficients of correlation are still significant but with lower values (r_5m_robust_ = 0.47; r_10m_robust_ = 0.62; r_15m_robust_ = 0.6).

