## Supplementary materials for "Mental Timing in Locomotion: Walking Across Real and Imagined Distances"

**Table S1**: Mean and standard deviations in seconds.

|  | Distance | | |
| --- | --- | --- | --- |
| Modality | 5m | 10m | 15m |
| Execution (seconds) | 3.71 ± 0.51 | 7.11 ± 0.83 | 10.51 ± 1.19 |
| Imagery (seconds) | 3.93 ± 1.03 | 7.35 ± 2.08 | 10.68 ± 3.13 |

**Table S2:** Bland-Altmann Test

| Distances | Limits | Values |
| --- | --- | --- |
| 5m | Upper Bound | 1.171 |
|  | Mean | -0.220 |
|  | Lower Bound | -1.611 |
| 10m | Upper Bound | 2.605 |
|  | Mean | -0.240 |
|  | Lower Bound | -3.085 |
| 15m | Upper Bound | 4.147 |
|  | Mean | -0.169 |
|  | Lower Bound | -4.485 |

**Table S3**: Correlations with MIQ-3 sub-scores

| Modality | | | | | |
| --- | --- | --- | --- | --- | --- |
|  | Kinesthetic | Visual 1st | Visual 3rd | K+V1 | Global |
| r | -0.201 | 0.014 | -0.232 | -0.106 | -0.233 |
| p-value | 0.287 | 0.942 | 0.218 | 0.576 | 0.216 |

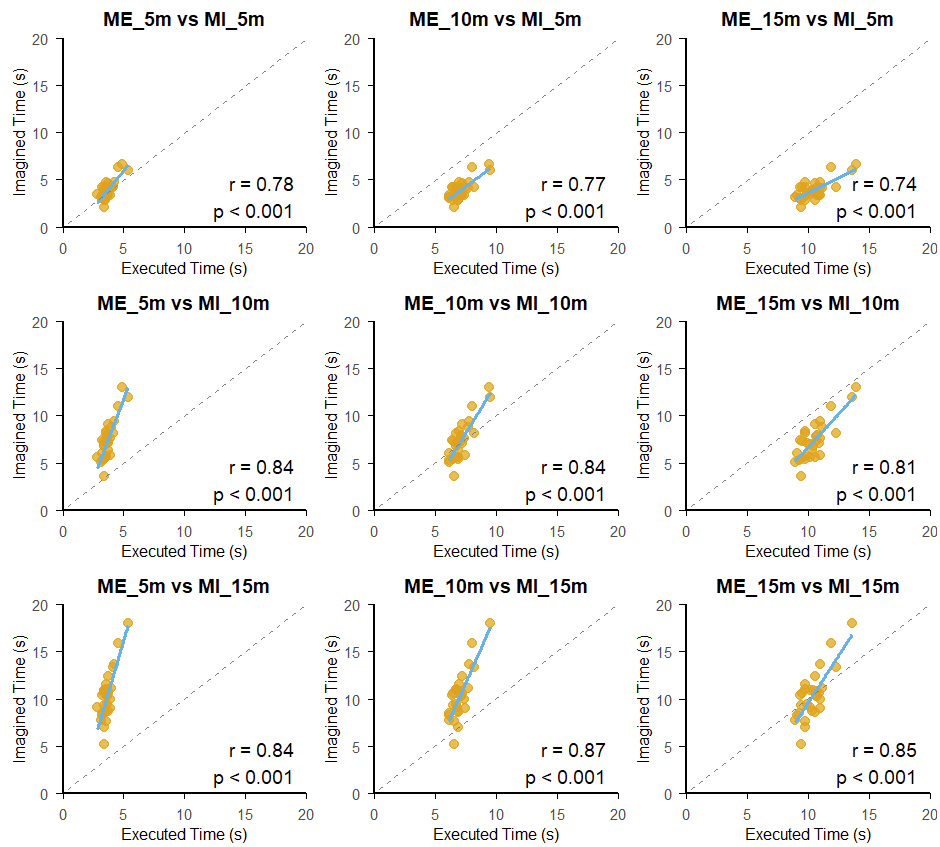

**Figure 1S:** Correlations across all distances and all modalities.

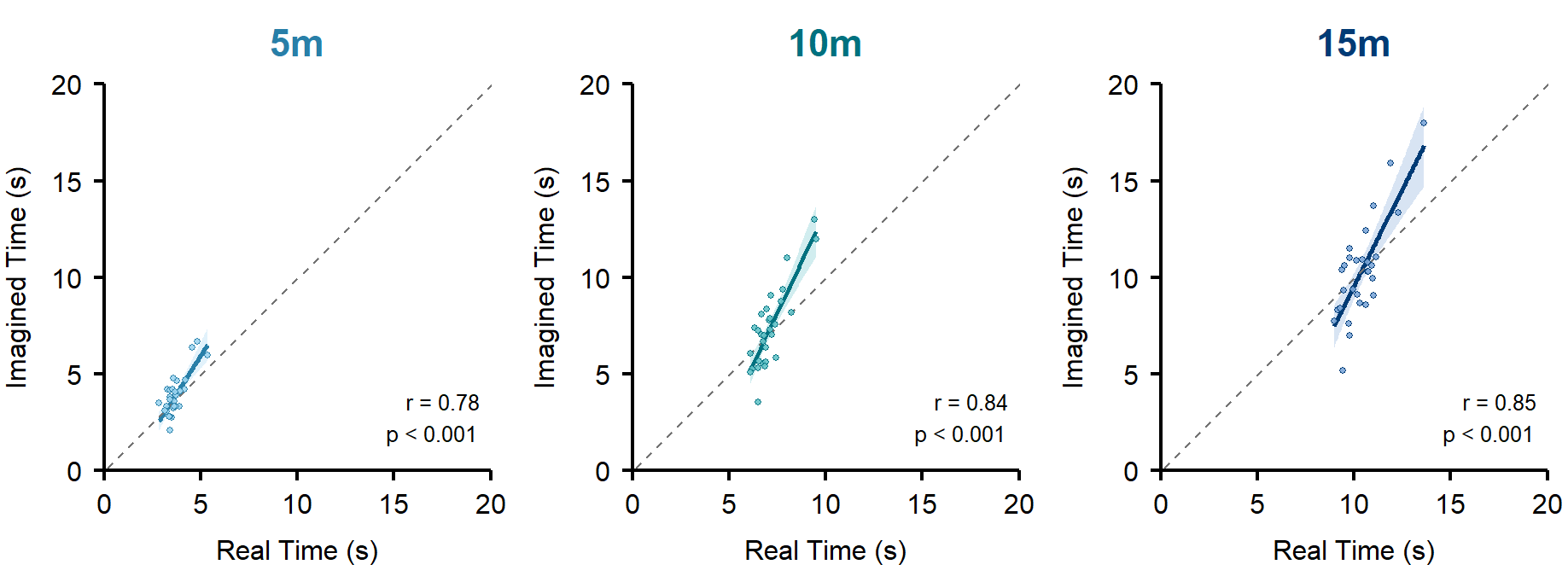

**Figure 2S:** Robust correlations. The points circled in purple are the outliers detected by robust correlation analysis. By removing these points, coefficients of correlation are still significant but with lower values (r_5m_robust_ = 0.47; r_10m_robust_ = 0.62; r_15m_robust_ = 0.6).
